# Intraluminal pressure triggers a rapid and persistent reinforcement of endothelial barriers

**DOI:** 10.1101/2025.01.22.634268

**Authors:** Aurélien Bancaud, Tadaaki Nakajima, Jun-Ichi Suehiro, Baptiste Alric, Florent Morfoisse, Jean Cacheux, Yukiko T. Matsunaga

## Abstract

In response to mechanical cues, endothelial cells elicit highly sensitive cellular response pathways that contribute to the regulation of the physiology and disorders of the vascular system. However, it remains relatively unexplored how endothelial tissues process and integrate the intraluminal pressure, and in turn regulate the permeation flow across the vessel wall. Leveraging a tissue engineering approach to create microvessels (MVs), we measured real-time permeation flow induced by intraluminal pressures ranging from 0.1 to 2.0 kPa. Our findings reveal that mechanically stimulated MVs strengthen their barrier function within seconds of exposure to pressures below 1 kPa, with this enhanced barrier function persisting for 30 minutes. We demonstrate that this barrier reinforcement is linked to the closure of paracellular gaps. Additionally, we observe that it is associated with, and depends on, actin cytoskeleton reorganization, including the accumulation of stress fibers near intercellular junctions and the broadening of adherence junction protein localization. These findings provide insights into the ability of endothelial tissues to regulate interstitial fluid flow in response to sudden increases in blood pressure.

---

Normal organ function relies on the precise regulation of blood vessel permeability, which governs the influx of molecular and macromolecular components from plasma (*1*). This barrier function is actively maintained in quiescent vasculature (*2*), primarily through organ-specific regulation of intercellular junction tightness (*3*). It is also dynamically responding to biochemical stimulations, e.g. thrombin (*4*) or histamine (*5*), that destabilize the endothelial barrier function (*6*) by disconnecting junctions and gradually forming intercellular separations. This destabilization is also associated to the reorganization of the actin cytoskeleton (*7*), a hallmark of mechanotransduction (*8*), wherein biochemical signals induce structural and physical changes within endothelial cells. Conversely, mechanotransduction triggered by shear stress (*9*, *10*) prompts elongation and alignment of endothelial cells in the direction of flow (*11*) while enhancing barrier integrity under shear stress below ∼1 Pa (*12*). Similar effects are observed in cyclic uniaxial stretch experiments (*13–16*), where elongating endothelial tissues induce cytoskeletal reorganization perpendicular to the strain direction and strengthen the barrier function at strains below ∼5%. At the molecular level, these adaptations involve the junctional mechanosensory complex (*17*), which orchestrates stress fiber assembly and regulates the distribution of junctional proteins (*18*, *19*). While these mechanisms have been extensively studied using fluid shear stress as a mechanical cue and monitoring barrier integrity through electrical resistance-based methods (*20*), the dynamic permeability changes of stretched endothelial cell monolayers remain more elusive. Circumferential stress is nevertheless a physiological cue of blood flow, which is characterized by a typical intraluminal pressure of ∼2 kPa in microvasculatures (*21*) with abrupt increases observed during hypertension (*22*).

To address this knowledge gap, we developed a technology that enables real-time monitoring of permeation flows across stretched endothelial cell monolayers. This technology is based on a microvessel-on-a-chip (MV) platform (*23–25*), which allows dynamic regulation of intraluminal pressure within the range of 50 to 2000 Pa. Permeation flow rates are simultaneously recorded over time windows spanning minutes to hours using a time interval of 100 ms. We demonstrate that the endothelial barrier actively responds to intraluminal pressure, becoming significantly less permeable within tens of seconds. We show that this reinforcement of the barrier is explained by the closure of paracellular pores, and report that these closure events can be challenged by an excessive intraluminal pressure of 1500 Pa or more. In the low-pressure regime of barrier reinforcement, we then prove that intraluminal pressure stimulation induces and depends on the reorganization of the actin cytoskeleton with the accumulation of stress fibers at the vicinity of intercellular junctions. Barrier reinforcement triggers, but is independent of, the activation of the MAP kinase mechanosensory pathway. Conversely, inhibition of the rho kinase compromises barrier reinforcement, destabilizing intercellular junctions for low intraluminal pressures. We finally discuss these findings in the context of endothelial tissue response to bursts in blood pressure.

## Results

### MVs constitute dynamic barriers to trans-endothelial flow

Using Human Umbilical Vein Endothelial Cells (HUVECs), which represent a standard for *in vitro* studies of vasculature and angiogenesis (*26*, *27*) and for endothelial phenotypic response to mechanical stimulation (*25*, *28*, *29*), we formed MVs of 200 µm in diameter and 6 mm in length within a collagen gel scaffold (Fig. 1A; detailed fabrication methods in the Methods section). After two days of culture in standard “static” conditions, we mounted a fluidic device onto the MV chip (Fig. 1B), enabling dynamic control of inlet pressures. By applying equal pressures to both inlets, we generated homogeneous intraluminal pressure without inducing fluid shear stress. The fluidic device and pressure actuation approach were adapted from previously reported methods for collagen gel mechanical characterization (*30*). To quantify the amount of fluid injected into the MV, flow meters were installed along the injection tubing (Fig. 1B). This setup allowed us to stimulate the MV with intraluminal pressure and simultaneously measure the permeation flow rate (𝑄) across the endothelial tissue and collagen matrix (indicated by red arrows in Fig. 1B). Additionally, the system’s integration with a wide-field microscope enabled cumulative observation of MV deformation over time (Fig. 1C-D).

**Figure 1:**
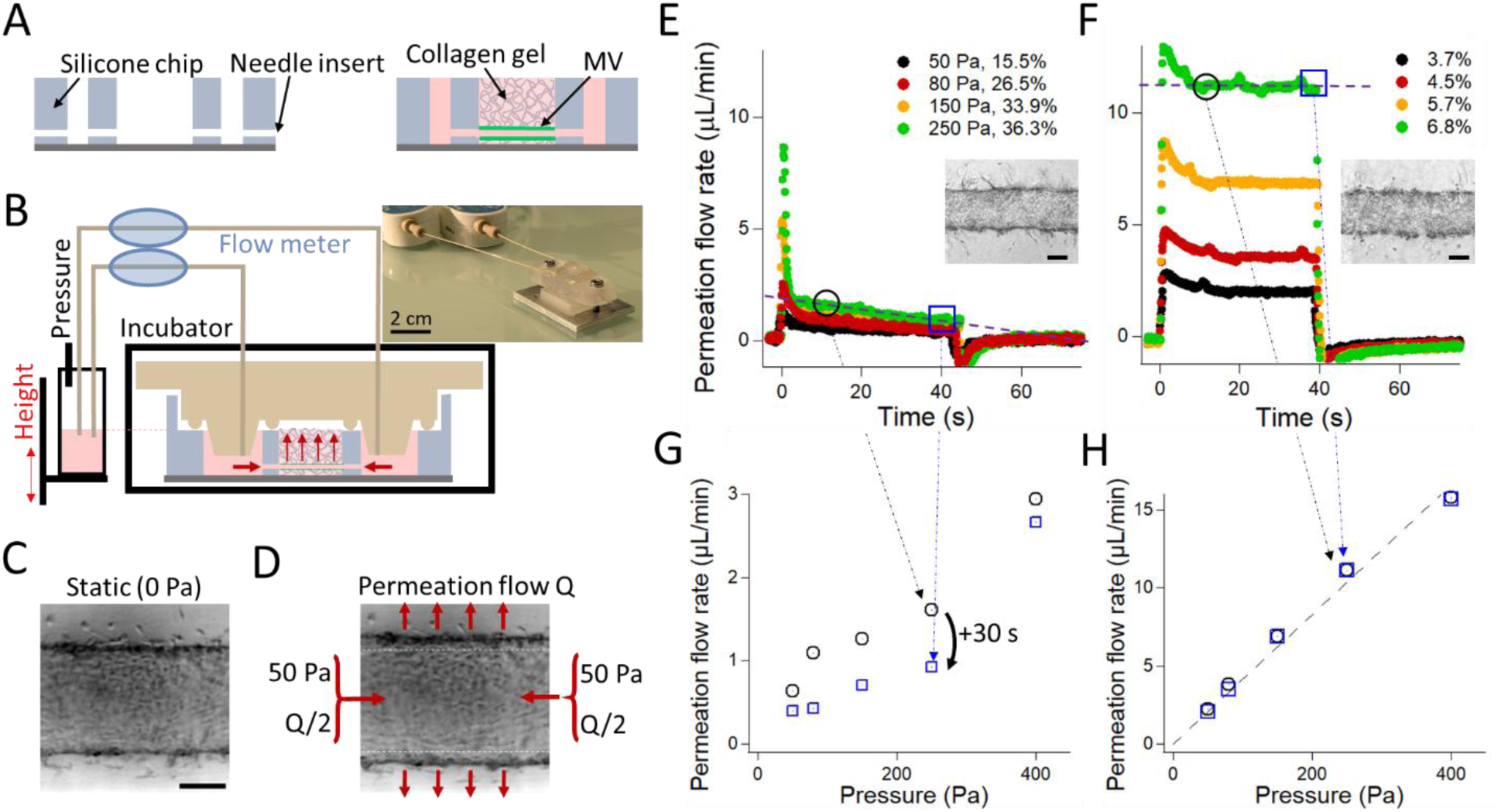
Real-time recording of the permeation flow through MVs. **(A)** The sketch represents the silicone chip to fabricate a hollow lumen in a collagen gel, which is seeded with endothelial cells to form the MV (green lines). **(B)** Schematics of the system to apply intraluminal pressure and record the permeation flow rate with a photograph. **(C)** The optical micrograph shows the MV in static conditions. **(D)** The same MV as in (C) upon application of an intraluminal pressure of 50 Pa. **(E)** The graph shows the permeation flow rate as a function of time for an intraluminal pressure spanning 50 to 250 Pa, as indicated in the legend. **(F)** The permeation flow rate is recorded after overnight treatment with sodium azide of the same MV as in (E). The micrographs in (E) and (F) show the MV and the lumen with cell residues, respectively. **(G)** The graph represents the flow rate as a function of the intraluminal pressure, as measured after 10 and 40 s of actuation (black and blue datasets, respectively). **(H)** Same as (G) after sodium azide treatment. The scale bars correspond to 100 µm.

We started by applying intraluminal pressures spanning 50 to 400 Pa, which typically compared to the stress induced by the pulsatile pressure in human capillaries of ∼300 Pa, but not to its continuous component of 2000 Pa (*31*). The resulting permeation flow traversing the MV was on the order of 𝑄 ∼1 µL/min, and the deformation of the MV in the range of 10 to 30% (Fig. 1E). The barrier function of the MV became evident when it was disrupted using the cytotoxic agent sodium azide, which caused a dramatic increase in flow through the collagen matrix—over tenfold for an intraluminal pressure of 250 Pa (Fig. 1F). In turn, these experiments enabled us to estimate the permeability of the MV, quantified by the Darcy coefficient (*32*). Considering the known permeability of collagen gels (7 10^-14^ m^2^, (*30*)) and the thickness of the collagen slab (2 mm) relative to the much thinner endothelial MV (∼1 µm, (*25*)), the observed tenfold reduction in permeation flow suggested an endothelial tissue permeability of approximately 5 10^-18^ m^2^ (see the details of the calculation in the methods section). This measurement was in agreement with published estimates (*25*, *32*), which modeled trans-endothelial fluid transport as convection through paracellular holes. To confirm that fluid transport occurred primarily via paracellular, rather than transcellular, pathways (*33*), we fixed the MV with formaldehyde immediately after manipulation. The permeation flow rate 𝑄 remained similar in amplitude before and after the fixation (Supplementary Fig. S1A), indicating that fluid transport was passive and occurred through paracellular pores, rather than being actively mediated by endothelial cells.

We then investigated the temporal variation of the permeation flow rate. In experiments with dead cells (Fig. 1F), we observed a rapid reduction in the flow rate during the first 10 seconds, which was attributed to the poroelastic deformation of the lumen (Supplementary Fig. S1B). Following this transient phase, the flow rate stabilized (dashed line in Fig. 1H), consistent with the behavior of passive porous materials such as collagen gels. In contrast, experiments with MVs exhibited a gradual decline in the flow rate over time (dashed line in Fig. 1E). To quantify this trend, we plotted the flow rate measured at 10 and 40 s that decreased across all pressure settings (Fig. 1G). This progressive decline in flow rate during stimulation suggests a change in the permeability of the endothelial cell monolayer, likely reflecting its active response to the mechanical cue.

### Intraluminal pressure stably reinforces MV barrier function

We further characterized the active response of the stimulated MV to determine whether the modulation of the permeation flow rate by intraluminal pressure was transient or stable. To investigate this, we applied an intraluminal pressure of 800 Pa for 150 s, released the pressure for 60 s, and then re-applied the stimulation for an additional 50 s. The pressure is shown in the black dataset, with the y-axis on the right in Fig. 2A, and the resulting permeation flow rate in the green dataset on the left y-axis. This experiment confirmed the decrease of 𝑄, which started from 2.0 and ended up at 1.1 µL/min in the first step of this experiment. The reduction was consistently observed in 24 MVs stimulated by an intraluminal pressure of 400 or 800 Pa and equal to -36 +/- 20% (Fig. 2B). On the contrary, it was significantly lower and equal to -7 +/- 2% in 14 controls obtained either by disrupting the tissue with sodium azide or by chemical fixation with paraformaldehyde (Fig. 2B). Then, 𝑄 returned to its initial null value as we stopped the pressure stimulation (Fig. 2A). More interesting was the fact that 𝑄 remained low during the second pressure pulse, with a value that precisely matched that at the end of the first stimulation of 1.2 µL/min. Given that a decrease in permeation flow indicates an enhanced barrier, referred to here as ’reinforced’, our data suggested a sustained barrier reinforcement following pressure stimulation. This result was strengthened by monitoring the permeation flow during ∼30 minutes following an initial stimulation of five minutes at 400 Pa (Fig. 2C). Barrier reinforcement occurred within the first two minutes; however, we observed a slow relaxation phase over the ensuing 30 minutes while maintaining zero pressure and applying brief pulses of ∼7 s to register 𝑄 (dashed line in Fig. 2C). We concluded that MV barrier reinforcement was rapid and persistent, in agreement with the swift activation of mechanosensitive pathways by fluid shear stress and the slow return to baseline (*9*, *10*).

**Figure 2:**
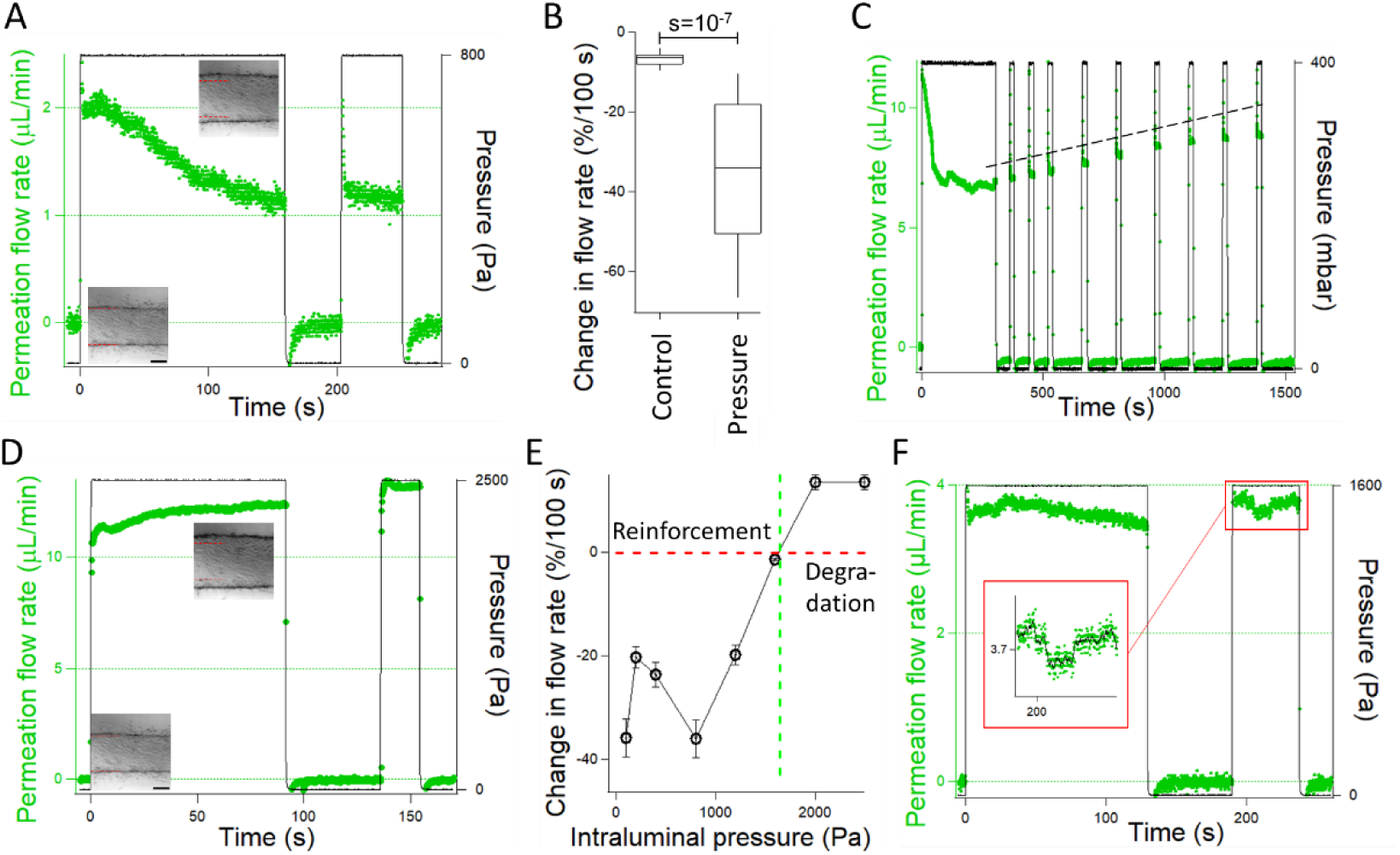
Rapid and persistent barrier reinforcement under intraluminal pressure. **(A)** The graph shows the permeation flow rate as function of time (green dataset) using a temporal sequence of pressure actuation represented in black and reported on the right axis. The two micrographs represent the MV in static and deformed by intraluminal pressure, at the bottom and top of the graph, respectively. **(B)** The box plot reports the relative change in permeation flow rate in the low-pressure regime in live MVs or control based on fixed or dead cells. **(C)** Same experimental design as in (A) but the temporal sequence of ∼30 min is ten times longer. **(D)** Using the same MV as in (A), we apply an intraluminal pressure of 2500 Pa. **(E)** The graph reports the relative change in permeation flow rate after a time lag of 100 s as a function of the intraluminal pressure. The red dashed curve corresponds to a constant permeation flow over time. The green dashed line draws the limit between the regimes of reinforcement and degradation. **(F)** In the intermediate pressure regime of 1600 Pa, the permeation flow rate is recorded over time. The scale bars correspond to 100 µm.

High tensile forces can disrupt adhesion between endothelial cells (*34*). Therefore, we anticipated that applying excessive intraluminal stress would destabilize the barrier. We repeated the experiment with the same MV as in Fig. 2A, but increased the intraluminal pressure to 2,500 Pa (Fig. 2D). The permeation flow rate was observed to increase during mechanical stimulation, and the endothelial tissue remained degraded even after a 60-second pause. Structural analysis of the MVs using immunoconfocal microscopy confirmed that the stress-induced barrier degradation resulted in detectable gaps between adjacent cells (Supplementary Fig. S2). By scoring the relative change in permeation flow rate during 100 s, we could then recapitulate a set of experiments performed on the same MV with two regimes of barrier reinforcement and barrier degradation (Fig. 2E). At the transition for an intraluminal pressure of 1,500 Pa, the permeation flow rate remained roughly stable over time (Fig. 2F). However, a closer inspection of the signal revealed abrupt positive and negative fluctuations of 𝑄. Using a Bayesian estimator to detect abrupt changes in temporal traces (*35*), we registered a total of 35 jumps for a total recording time of 500 s and 5 different MVs. Their average amplitude was 95 +/- 81 nL/min (positive and negative jumps being roughly equally frequent, Supplementary Fig. S3). At the threshold pressure, barrier reinforcement likely competed with the disruption of intercellular junctions, resulting in paracellular pore opening and closing events. Notably, this tentative mechanism could be used to estimate the typical size of these events. Indeed, the flow rate of ∼100 nL/min for a pressure of 1,500 Pa corresponded to a single pore approximately 1 µm in radius crossing an endothelial tissue of 1 µm in thickness (see the details of the calculation in the methods section). Consequently, analysis of the dynamics of barrier response to intraluminal pressure hints to a mechanism of reinforcement involving paracellular pore closure.

### Paracellular pore closure events account for the reinforcement of the barrier

We aimed to gather further evidence on whether barrier reinforcement involved paracellular hole closure. We performed the macromolecular assay, which consisted in loading a fluorescent dextran molecule into the lumen of the microvessel (MV) and tracking its spatial redistribution within the surrounding extracellular matrix (Fig. 3A). The experiment was notably performed under constant intraluminal pressure, maintained at 100 Pa throughout the assay, as described in refs. (*25*, *36*), starting two minutes prior to dye injection. Leakage through paracellular gaps became immediately detectable following dye injection (upper left panel in Fig. 3A), because these gaps created an escape path for the rapid release of fluorescent molecules within the basal layer of the MV. These bright spots gradually spread over time due to diffusion, as illustrated in the difference micrograph between consecutive images (upper panel in Fig. 3B). Interestingly, this differential representation also revealed the appearance of localized dark spots at 15 and 38 s (Fig. 3B). These dark spots indicated an abrupt cessation of fluorescence convection, as expected for the closure of paracellular pores. These closure events occurred only once, as exemplified by the observation that the dark spot appearing after 15 s became undetectable at 38 s. Although spatial analysis beyond one minute was not possible with this assay, because the global fluorescence signal induced by leakage blurred out local intensity fluctuations, these experiments provided direct supporting evidence of localized rearrangements of the endothelial tissue architecture that reduce the strength of the permeation flow.

**Figure 3:**
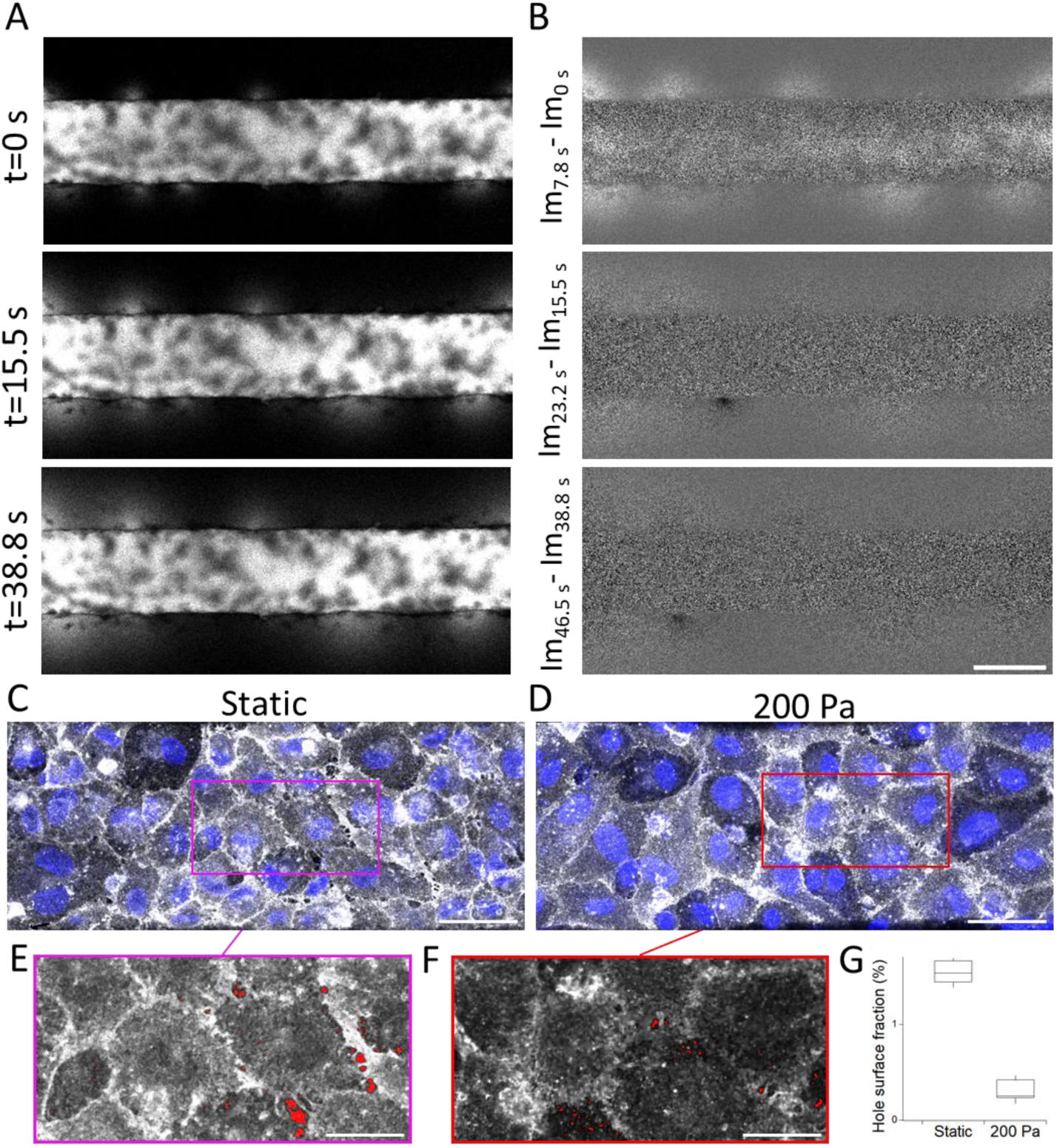
Paracellular gap closure occurs under pressure. **(A)** Confocal time series of the macromolecular assay using the 4 kDa dextran performed with an intraluminal pressure of 100 Pa using the system described in (36). The inter-frame interval is 7.8 s. **(B)** The micrographs correspond to the difference between two consecutive images of the time series in (A). **(C)** MIPs of confocal micrographs of a MV cultured in static conditions; labels: PECAM-1 in gray and DNA in blue. **(D)** Same as (C) after the application of an intraluminal pressure of 200 Pa. Note that the spatial distribution of PECAM-1 appears more diffuse, as discussed in the following section. **(E)** and **(F)** are zoom-ins extracted from panels (C) and (D) with the detected paracellular gaps in red. **(G)** Surface fraction of gaps in static and after pressure stimulation. The scale bars correspond to 200 µm in A-B, 50 µm in C-D, and 25 µm in E-F.

Second, we aimed to validate the presence of paracellular pores by characterizing the structure of the endothelial cell monolayer with immunoconfocal microscopy performed in static conditions *vs*. under intraluminal pressure. We used cell-cell junction protein platelet endothelial cell adhesion molecule-1 (PECAM-1) to assess intercellular interactions. Qualitative inspection of MVs revealed occasional gaps, indicated by dark fluorescence signals under static conditions (Fig. 3C). These gaps were less frequent following intraluminal pressure stimulation at 200 Pa for 5 minutes (Fig. 3D). Although the frequency of these gaps varied across fabrication batches, we estimated the gap surface fraction using image analysis (see Methods, Fig. 3E–F) across two separate fabrication batches, including a total of 4 MVs in static conditions and 6 under mechanical stimulation. The average area fraction of gaps was estimated to be 1.55 ± 0.12% in static conditions, compared to 0.31 ± 0.11% after stimulation (Fig. 3G). Altogether, our data strongly support the conclusion that endothelial barriers respond and adapt to intraluminal pressure by closing paracellular gaps.

### Intraluminal pressure stimulates actin cytoskeleton reorganization

The actin cytoskeleton plays a central role in regulating endothelial tissue function, as demonstrated by the morphological and orientational changes endothelial cells undergo in response to mechanical stimulation from fluid shear stress or periodic stretching (*17*, *37*). We thus characterized the actin cytoskeleton and the patterns of cell-cell interaction proteins in order to clarify their structure after barrier reinforcement. We stained HUVECs with phalloidin and antibodies to the *adherens* junction (AJ) protein Vascular Endothelial Cadherin (VE-Cad), PECAM-1 or the tight junction protein Claudin-5 (see Supplementary Fig. S4 for VE-Cad and Claudin 5). In the control samples cultured in static conditions, we observed a well-defined cortical actin rim at the basolateral site, indicating a typical cytoskeletal organization in the absence of pressure (Fig. 4A). We then stimulated MVs with 100 and 400 Pa of intraluminal pressure stimulation for 5 minutes, and fixed the samples immediately after. At 100 Pa, the strain was 17 ± 2%, and this pressure load was sufficient to induce the formation of parallel actin stress fibers spanning the cell interior (Fig. 4B and Supplementary Fig. S4A). In some instances, we observed alignment of stress fibers between neighboring cells, accompanied by irregular intercellular junction patterns, an arrangement reminiscent of discontinuous AJs ((*38*), orange arrows in Fig. 4B). Notably, these patterns closely resembled those observed upon stimulation of endothelial tissues with pro-inflammatory agents, such as tumor necrosis factor or histamine, both of which are associated with hyperpermeability (*5*, *39*, *40*). However, unlike these biochemical cues, mechanical stimulation-induced formation of aligned stress fibers and discontinuous AJs did not lead to a degradation of endothelial barrier function. Examination of stress fiber localization revealed their predominant accumulation on the apical surface of the tissue, with a few isolated fibers observed on the basal side (lower panel of Fig. 4B). When intraluminal pressure was increased to 400 Pa, actin stress fibers became concentrated near cell-cell junctions, as indicated by the orange-labeled regions in Fig. 4C. This pattern was also evident in VE-Cadherin MIPs (Supplementary Fig. S4A). Notably, at 400 Pa, actin stress fibers were exclusively observed on the apical side (Fig. 4C), and discontinuous AJ patterns appeared less frequently. However, the intercellular junctions visualized in MIPs were broader under 400 Pa compared to static conditions, as quantified by the onset of 29% of the full width at half maximum of the Gaussian fit to the PECAM-1 signal (Fig. 4D).

**Figure 4:**
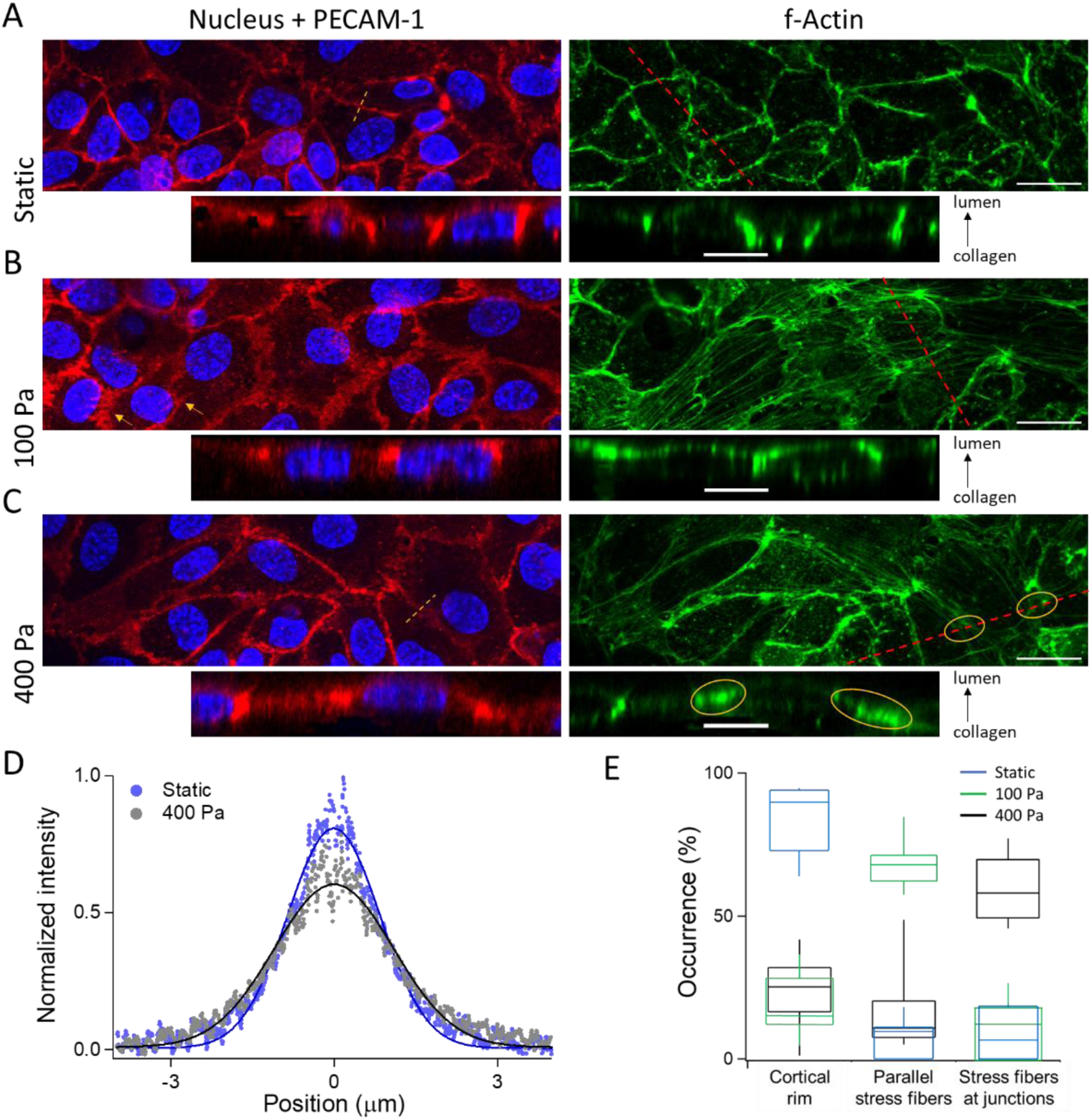
Barrier enhancement involves actin cytoskeleton reorganization. MIPs and vertical cross-sections along the red dashed lines in static conditions **(A)** and after the application of 100 Pa **(B)** or 400 Pa **(C)**; labels: f-Actin in green, DNA in blue, and PECAM-1 in red. The scale bars correspond to 20 µm in the MIPs and 10 µm in the cross-sections (see Supplementary Fig. S4 for VE-Cad data). **(D)** Integral-normalized intensity profile of the AJs extracted from MIP of PECAM-1 immunoconfocal micrographs. 30 intensity profiles, which typically correspond to the profile along the brown dashed lines in the panels (A) and (C), are pooled in each condition. **(E)** Enumeration of the number of HUVECs with a cortical rim, parallel stress fibers or accumulated stress fibers in static conditions or under pressure.

Based on these observations, we identified three distinct cytoskeletal patterns: (i) a cortical rim, (ii) parallel stress fibers, and (iii) accumulated stress fibers. These patterns were counted under static conditions and at pressures of 100 and 400 Pa (blue, green, and black datasets in Fig. 4E), with data averaged across three microvessels, including ∼120 cells, per condition. This analysis confirmed the phenotypic shift induced by intraluminal pressure because the prevalence of the three cytoskeletal patterns peaked at 84%, 55%, and 61% for the three different settings in intraluminal pressure. Thus, the intraluminal load stimulates cytoskeletal reorganization, which in turn stabilizes intercellular junctions.

Further, we tested whether barrier reinforcement was dependent on the mitogen-activated protein kinase (MAPK) pathway because cyclic-stretch experiments have shown that mechanical stretch activates this mechanosensory cascade independently of actin fiber reorientation (*41*). We used the selective MEK 1/2 inhibitor PD98059 at 30 µM and examined the response of MVs to intraluminal pressure. Immunostaining for the phosphorylated form of extracellular signal-regulated kinase (ERK) confirmed that the pressure stimulus activated the MAPK pathway and that this activation was inhibited by PD98059 (Supplementary Fig. S5A-B). We also monitored the permeation flow rate for 5 minutes with or without the inhibitor (Supplementary Fig. S5C), and did not detect any changes of the barrier reinforcement dynamics. Last, we observed that the density of holes in static conditions decreased similarly after pressure stimulation with or without drug treatment (Supplementary Fig. S5D). Consequently, the inhibitor did not affect the response of the endothelial barrier to intraluminal pressure, indicating that the activation of the MAPK pathway was coincident but not associated to the endothelial tissue reinforcement.

### Barrier reinforcement is lost upon inhibition of ROCK

The reorganization of the actin cytoskeleton occurred concurrently with endothelial adaptation to intraluminal pressure. To further investigate this process, we aimed to disrupt actin reorganization by inhibiting ROCK — a kinase activated by RhoA that regulates myosin motor activity and cell contractility — with Y27632. This inhibitor has been shown to directly interfere with actin stress fiber formation without disrupting endothelial barrier function at concentrations below 10 µM (*42*); at these low concentrations, it even protects the barrier against thrombin-induced degradation (*43*, *44*). After one day of culture in static conditions, we supplemented the medium with 1 µM of Y27632 and maintained the MVs for another two days. This treatment did not induce any morphological change at the tissue level (Fig. 5A), and it also preserved the characteristic cortical rim pattern in mechanically unstimulated MVs the (see more below). We then probed the response of MVs to intraluminal pressure by applying 400 Pa. The initial flow rate after a stimulation of ∼1 s was comparable in the control and in the treated sample of 7.6 and 7 µL/min, respectively (gray and green datasets in Fig. 5B). These results demonstrated that the permeability of the barrier was not affected by Y27632 treatment. However, the response diverged noticeably after approximately 10 seconds of actuation with intraluminal pressure. Specifically, the control sample exhibited a characteristic rapid and sustained reinforcement of the barrier function, as shown by the red dashed trend line in Fig. 5B. In contrast, the permeation flow rate in Y27632-treated MVs remained consistently high and unchanged, even after 50 seconds (blue dashed line in Fig. 5B). This flat response indicated that the endothelial barrier lost its ability to actively respond to intraluminal pressure following treatment with Y27632. This conclusion was further supported by measuring the change in permeation flow rate after 100 seconds of actuation. Unlike the untreated MVs (Fig. 2B), the Y27632-treated MVs showed no significant differences in response when compared to dead or fixed cells (Fig. 5C). Taken together, these findings suggest that treatment with 1 µM Y27632 compromises the adaptive reinforcement of endothelial tissue in response to intraluminal pressure.

**Figure 5:**
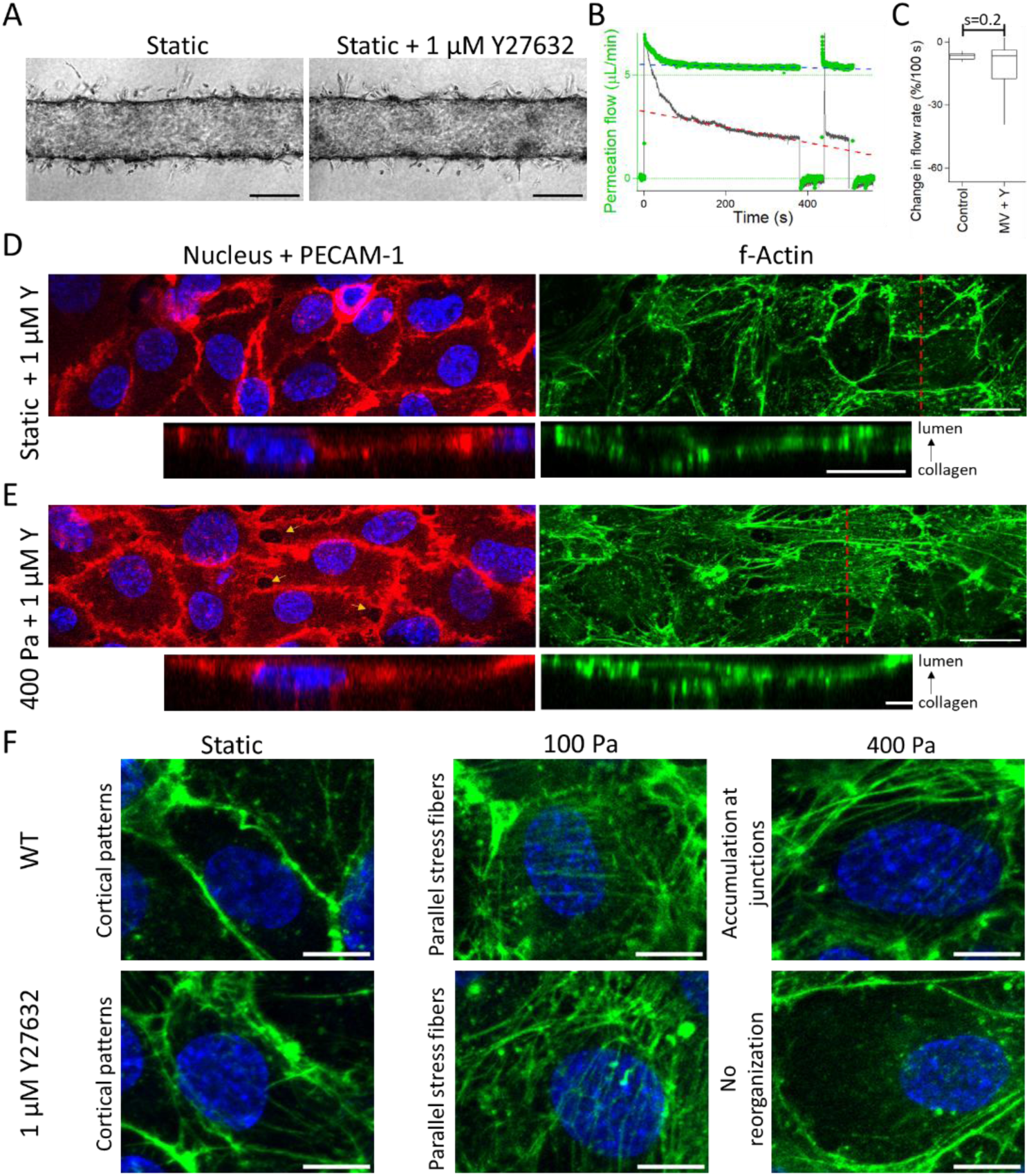
Rho kinase inhibition disrupts barrier reinforcement. **(A)** Optical micrographs of MV treated or not with the drug Y27632 in static conditions. **(B)** The graph shows the permeation flow rate as function of time for a MV treated with Y27632 (green dots) and untreated (gray dots). The temporal sequence of pressure actuation is the same as in Fig. 2A. **(C)** The box plot reports the relative change in permeation flow rate after 100 s of stimulation with fixed or dead cells (same dataset as in Fig. 2B) or in treated conditions. MIPs of MVs treated with Y27632 and vertical cross-sections along the red dashed lines in static conditions **(D)** and after the application of 400 Pa **(E)**; labels: f-Actin in green, DNA in blue, and PECAM-1 in red (see Supplementary Fig. S6 for VE-Cad data). The scale bars correspond to 20 µm in the MIPs and 10 µm in the cross-sections. **(F)** MIPs of the apical surface of HUVECs, with an integration depth of 3 µm, under control and treated conditions. Labels: f-Actin in green, DNA in blue; the scale bars correspond to 10 µm.

We next analyzed the actin cytoskeleton and PECAM-1 patterns in the presence of the ROCK inhibitor using immunoconfocal microscopy (Fig. 5D-E). Under static conditions, the actin cortical rim and linear AJs were visible in the MIPs (Fig. 5D), exhibiting patterns similar to the untreated control group (Fig. 4A). However, after exposure to 400 Pa of pressure, we observed significant alterations compared to the control. Specifically, discontinuous actin fibers were detected (Fig. 5E), predominantly distributed along both the basal and apical surfaces of the cells. The AJ patterns appeared irregular, with protrusions extending along typically thin actin fiber tracks (Supplementary Fig. S6). Additionally, ovoid gaps between adjacent cells were also detected, suggesting that intraluminal pressure disrupted intercellular junctions (highlighted by orange arrows, Fig. 5E) and contributed to the loss of barrier integrity under treated conditions. In order to further document the loss of cytoskeleton reorganization induced by Y27632, we extracted the actin pattern on the apical side of the tissue in static conditions and after 100 and 400 Pa of pressure stimulation (Fig. 5F). While the actin patterns under static conditions and at 100 Pa were comparable between the control and treated groups, the reorganization and accumulation of stress fibers at junctions observed at 400 Pa in controls was completely absent in Y27632-treated MVs. These findings collectively indicate that MVs with impaired cytoskeletal reorganization cannot effectively withstand intraluminal stress, resulting in the formation of paracellular gaps and a compromised barrier function.

## Conclusion

Using an assay to dynamically measure the permeation flow across endothelial cell monolayers, we report that HUVEC-based MVs respond to intraluminal pressure stimulation in seconds by reinforcing their barrier function. We show that this reinforcement is explained by the active closure of paracellular holes. We also demonstrate that this adaptative response can be challenged by intraluminal pressures greater than 1,500 Pa, and uncover a dynamic equilibrium of opening and closing events at this threshold. Although the vasculature model used in this study is simplified regarding the extracellular matrix protein composition and perivascular cell integration, we note that this threshold is comparable to the physiological blood pressure in capillaries, approximately 2000 Pa (*45*). This finding suggests that endothelial cells alone have a strong capacity to withstand the constraints imposed by blood pressure.

Our data further highlight the crucial role of actin cytoskeleton reorganization in reinforcing barrier function under mechanical stress. At a low intraluminal stress of 100 Pa, we observed parallel actin stress fibers spanning the cell interior, accompanied by zig-zag patterns of cell-cell adhesion proteins. These patterns resemble the perpendicular stress fibers at AJs commonly associated with inflammatory conditions (*38*). While this cytoskeletal organization may be prone to destabilization, it undergoes a significant rearrangement at 400 Pa, where parallel actin fibers localize at cell junctions. Given that intercellular junctions in HUVECs are stabilized by the pushing forces of the branched actin network emerging from cortical actin bundles (*46*), we propose that the accumulation of actin stress fibers at junctions enhances cell-cell association forces. This process effectively seals potential gaps in the tissue. Interestingly, this model suggests a counterintuitive mechanism in which increased intraluminal pressure, which stretches the tissue, triggers a corresponding rise in compressive tension between cells, thereby strengthening the barrier.

Finally, it is tempting to propose that the reinforcement of endothelial barriers plays a key role in regulating interstitial fluid flow and preventing the onset of edema. Increases in intraluminal pressure, commonly seen in hypertension or during physical exercise (*47*), raise the pressure within blood vessels, including capillaries. When capillary pressure rises, permeation flow also increases, heightening the risk of fluid accumulation in tissues. Prolonged hypertension has been shown to induce structural changes in blood vessels, such as vessel wall thickening, which helps resist elevated pressure and limit interstitial fluid leakage (*48*, *49*). Aside from long-term vascular remodeling, barrier reinforcement offers a rapid and durable mechanism for regulating permeation flow, potentially maintaining tissue homeostasis despite sudden changes in blood pressure.

## Materials and methods

### Cell culture and reagents

All chemicals were purchased from Sigma Aldrich, unless mentioned. Studies were performed with Primary human umbilical vein endothelial cells (HUVEC; Catalog #C2519A, Lot #0000699241; Lonza, Basel, Switzerland) that were cultured in Endothelial Cell Growth Medium-2 BulletKit (EGM-2; Lonza). They were frozen in liquid nitrogen at passage 4 to 5, thawed and cultured for three days in culture dishes, and then used to load MV chips. The drug PD98059 was dissolved at 50 mM in anhydrous DMSO and diluted in culture medium to 30 µM. Y27632 was purchased at 1 mM in H20 and diluted in culture medium at 1 µM.

### MV fabrication

MV were fabricated in polydimethylsiloxane (PDMS)-based chips (25 mm × 25 mm × 5 mm: width × length × height), as developed by Matsunaga and collaborators (*36*). The protocol includes an additional PDMS-collagen cross-linking step to avoid leaks at the PDMS/collagen interface during diffusion and pressure assays. The protocol started by O2 plasma treatment of PDMS chips and acupuncture needles of 200 µm (No. 08, J type; Seirin, Shizuoka, Japan) for one minute (basic plasma cleaner; Harrick Plasma, Ithaca, NY, USA). The PDMS chips and needles were then placed together in a vacuum reactor with 50 µL of aminopropyl-triethoxysilane, and left at 0.1 mbar and room temperature for 30 minutes. Needles were then soaked in 1% (w/v) bovine serum albumin and dried. The chips were treated with 50 µL of 2.5% glutaraldehyde for one minute, then thoroughly rinsed with water and dried. The collagen solution was subsequently prepared on ice by mixing Cellmatrix® Type I-A collagen solution (Nitta Gelatin, Japan), 10× Hanks’ buffer, and 10× collagen buffer (volume ratio 8:1:1) following manufacturer’s protocol (final collagen concentration: 2.4 mg/mL). We poured 30 µL of this ice-cold solution into the chip, and inserted the coated needle. The resulting devices were incubated at 37°C for 15 min to induce collagen reticulation, and the needles were withdrawn to form a hollow channel. The chips were left in PBS at least overnight before cell seeding, and the holes for needle incorporation were sealed with unreticulated PDMS to prevent leaks.

Just prior to loading in the chips, HUVEC cells were harvested and resuspended in the medium supplemented with 3% (m/v) dextran (500 kDa) at a density of 10^7^ cells/mL. 50,000 cells were loaded at each opening of the channel, and let to attach to the collagen scaffold at 37°C for 10 minutes. We eventually added 1 mL of fresh medium, and refreshed it every other day.

### Microfluidics to monitor the permeation flow

MV were placed on an aluminum support of 30 × 60 mm^2^ with a set of tapped holes that enabled us to tightly hold 3D printed reservoirs fabricated by stereolithography (Expert Material Series, NSS, Japan). This system was connected to chromatography PEEK tubing of outer diameter 1/32” and inner diameter 380 µm. They were connected to flow sensors (Fluigent, model S) through UNF 6-40 port, and then to pressurized reservoirs of 2 mL actuated with a 25-mbar pressure controller (MFCS, Fluigent). These reservoirs were mounted on a manually adjustable vertical stage, allowing precise control of fluid levels to eliminate hydrostatic flows. Prior to every experiment, the microfluidic system was first filled with 70% ethanol solutions, then extensively washed with sterile medium, before its assembly on the MV chip. The mounted MV chip was eventually placed in a miniature incubator controlled in temperature and CO2 that was placed on an inverted bright field microscope (Zeiss AxioObserver). Note that the hydraulic resistance of the microfluidic set up of 15 Pa/(µL/min) was three and thirty times smaller than that of collagen gels and MV, respectively.

### Fixation and immunostaining

MV were fixed using 4% (w/w) paraformaldehyde for 30 min, and then thoroughly rinsed with PBS. Permeabilization was performed with 0.5% Triton X-100 for 10 min at room temperature. Blocking with 1% BSA was performed overnight at 4°C. Cells were then incubated overnight at 4°C with the primary antibodies VE-Cadh (rabbit mAb, D87F2, Cell Signaling Technology, 1:200), PECAM-1 (mouse mAb, GA610, Dako, 1:200), Claudin-5 (rabbit mAb, ab15106, Abcam, 1:200) or phosphorylated ERK1/2 (rabbit mAb, 4370T, Cell Signaling Technology, 1:200) diluted in blocking solution. After washing, cells were incubated for 2 h with the corresponding secondary antibodies (1:400), Alexa Fluor-488 Phalloidin (1:800), and Hoechst 33342 (1:1000). Labeled samples were washed and stored at 4°C until imaging. Confocal images were captured with the LSM 700 confocal microscope (Carl Zeiss) equipped with a 40× water immersion objective (numerical aperture of 1.2). We used a pinhole of 1 Airy unit for the three lasers of 405, 488, 555 nm, a pixel size of 0.156 µm, and set the vertical increment of confocal stacks to 0.5 μm.

### Image analysis and statistical test

Image analysis, including MIP and orthogonal view, was performed with ImageJ. Hole detection was performed by thresholding images (Minimum method) after application of the “enhanced local contrast” function (CLAHE). All the plots were obtained with Igor Pro (version 5.0). Statistical significance was determined using the Student’s t-test, that was reported with the parameter *s* (s-values < 0.05 were considered statistically significant; n.d., not different). Data were expressed as mean ± standard error.

### Qualitative estimation of MV permeability

The permeation flow velocity is related to the pressure gradient by the following relationship:

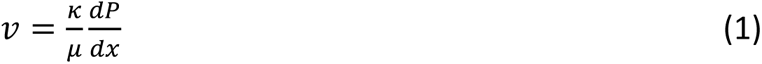

with 𝜅 the permeability and 𝜇 the viscosity of the culture medium. We assume a 1D geometry and a linear pressure gradient, implying that the pressure gradient is 𝑃_𝑖𝑛_⁄𝐻_𝑐_ with 𝐻_𝑐_ the collagen gel height of 1 mm. The endothelial barrier confines the pressure gradient in the cell layer ℎ. Using the subscripts *c* and *e* for the collagen gel and the endothelial cell layer, respectively, we deduce that:

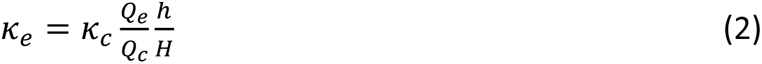

---

**Supplementary Figure S1:**
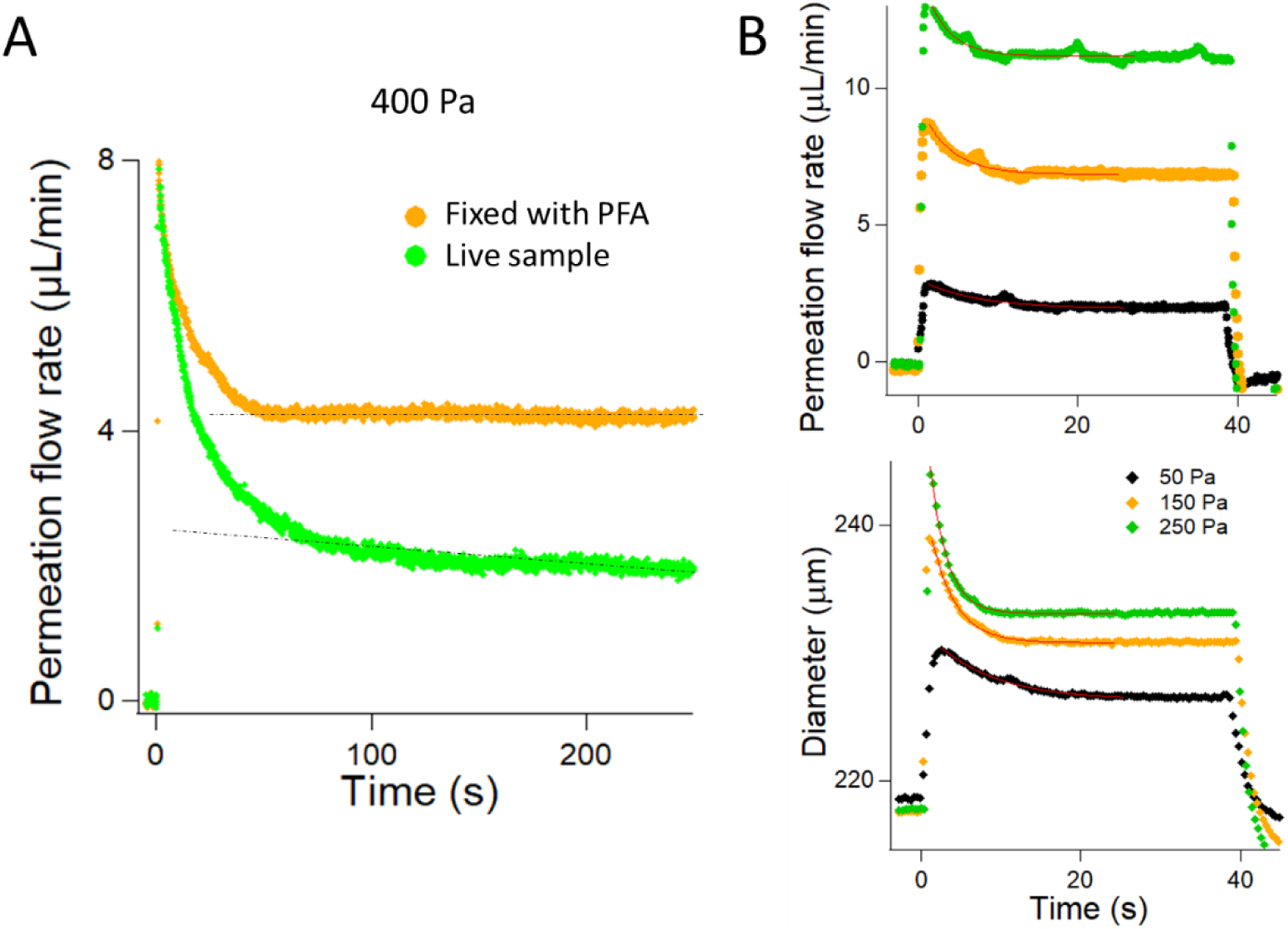
Characterization of the permeation flow and MV deformation. **(A)** Permeation flow rate as a function of time for the same MV live or fixed. The initial value of the permeation is comparable, and the reinforcement of the barrier is lost after fixation. **(B)** The flow rate through the collagen slab is shown in the upper panel for two different intraluminal pressure stimulation of 50 and 250 Pa. The deformation of the tube is represented in the lower panel. The transient regimes of deformation and permeation are closely correlated, as evidenced by the mono-exponential fitting of the deformation relaxation rates, which are 7.7 ± 0.4, 3.03 ± 0.09, and 2.13 ± 0.04 s, compared to the corresponding flow rate relaxation rates of 7.1 ± 0.4, 3.70 ± 0.15, and 2.70 ± 0.08 s for intraluminal pressure stimulations of 50, 150, and 250 Pa, respectively. This correlation arises from the deformation of the tube being linked to a transient regime of additional fluid injection, commonly referred to as the capacitance effect in microfluidics (*50*).

**Supplementary Fig. S2:**
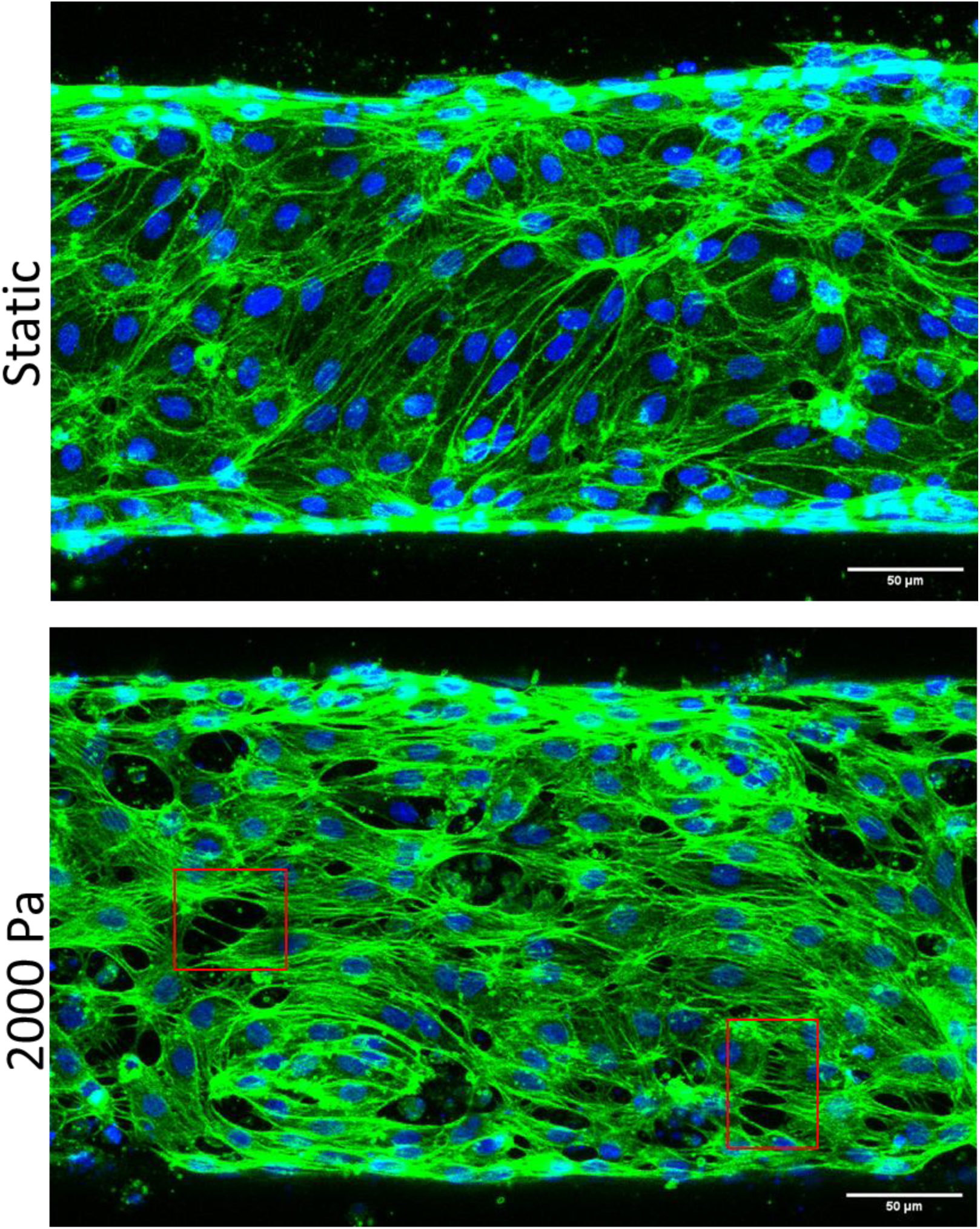
Structural inspection of MVs after stimulation with an intraluminal pressure of 2000 Pa. The upper panel is the MIP in static conditions and the lower panel presents the stressed MVs. The red rectangles present paracellular holes in the tissue. labels: f-Actin in green, DNA in blue. The scale bars correspond to 50 µm.

**Supplementary Fig. S3:**
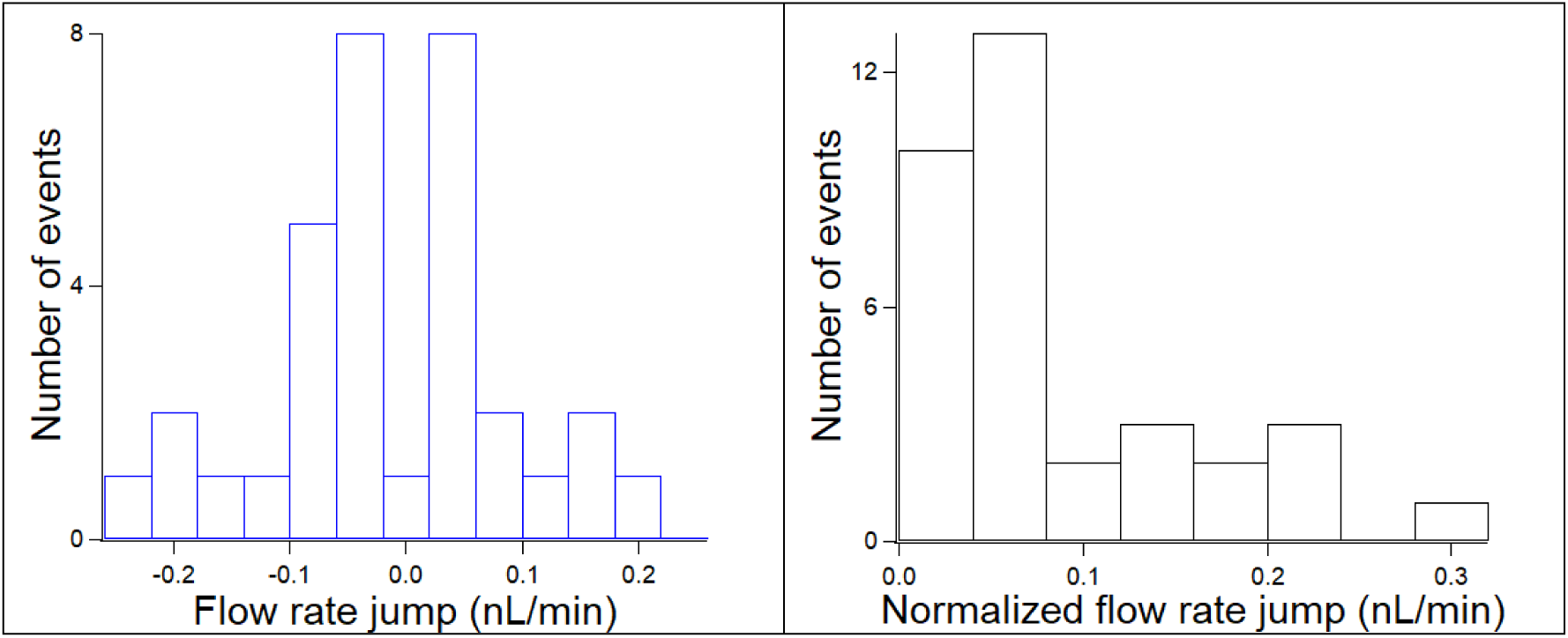
Analysis of the flow rate jumps at 1500 Pa. The left panel shows the histogram of the flow rate jumps. Note that the probability of positive and negative jumps is roughly equal. The histogram at the right is the same data after normalizing the jumps.

**Supplementary Fig. S4:**
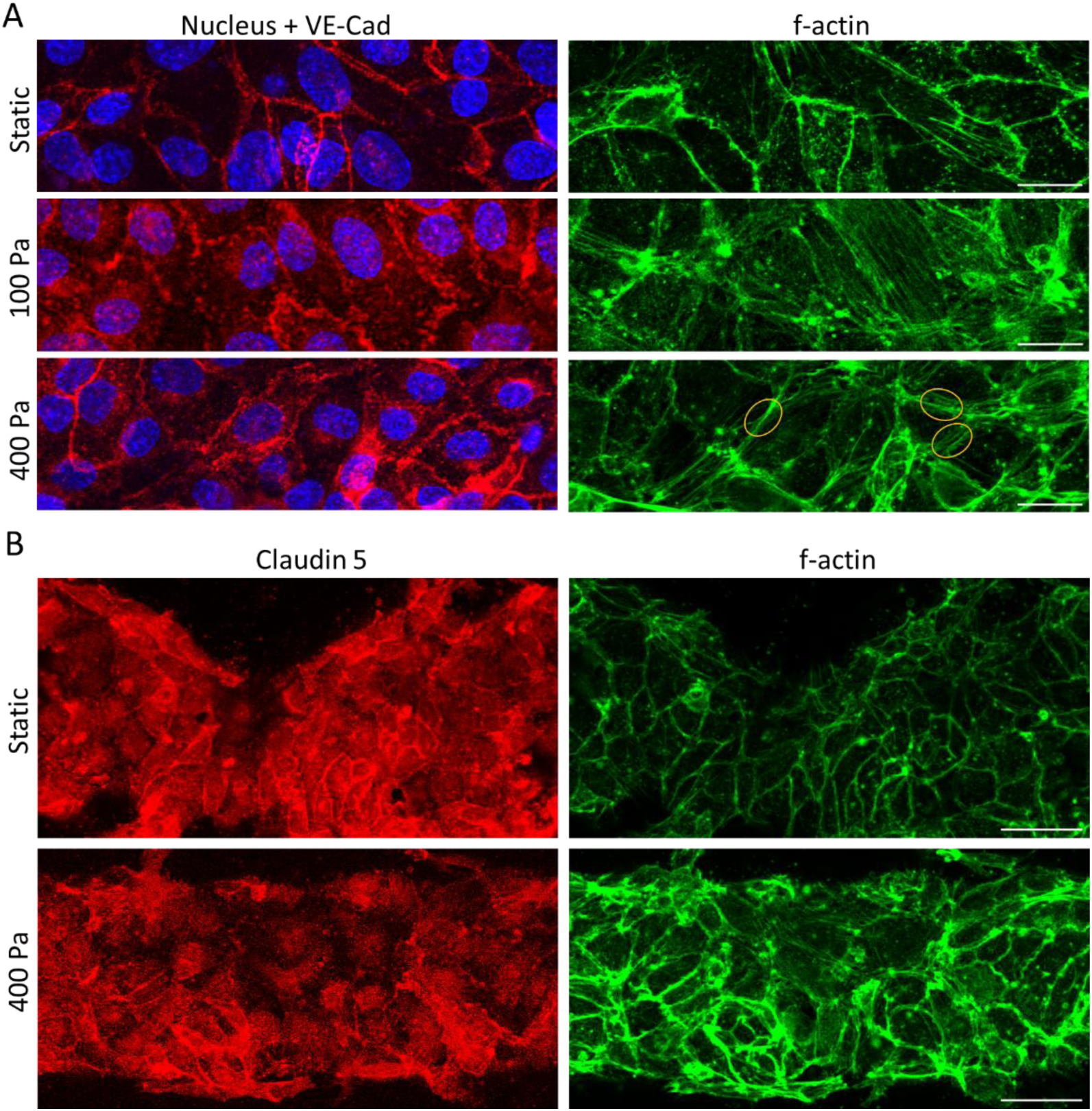
MIP of VE-Cad and Claudin in static or after pressure stimulation. **(A)** MIPs in static conditions and after the application of 100 or 400 Pa in the same imaging conditions as in main text; labels: f-Actin in green, DNA in blue, and VE-Cad in red. The scale bars correspond to 20 µm. **(B)** MIPs in static conditions and after the application of 400 Pa using a pixel size of 0.313 µm and a vertical inter-frame interval of 2.5 µm; labels: f-Actin in green, DNA in blue, and Claudin in red. The scale bars correspond to 50 µm.

**Supplementary Fig. S5:**
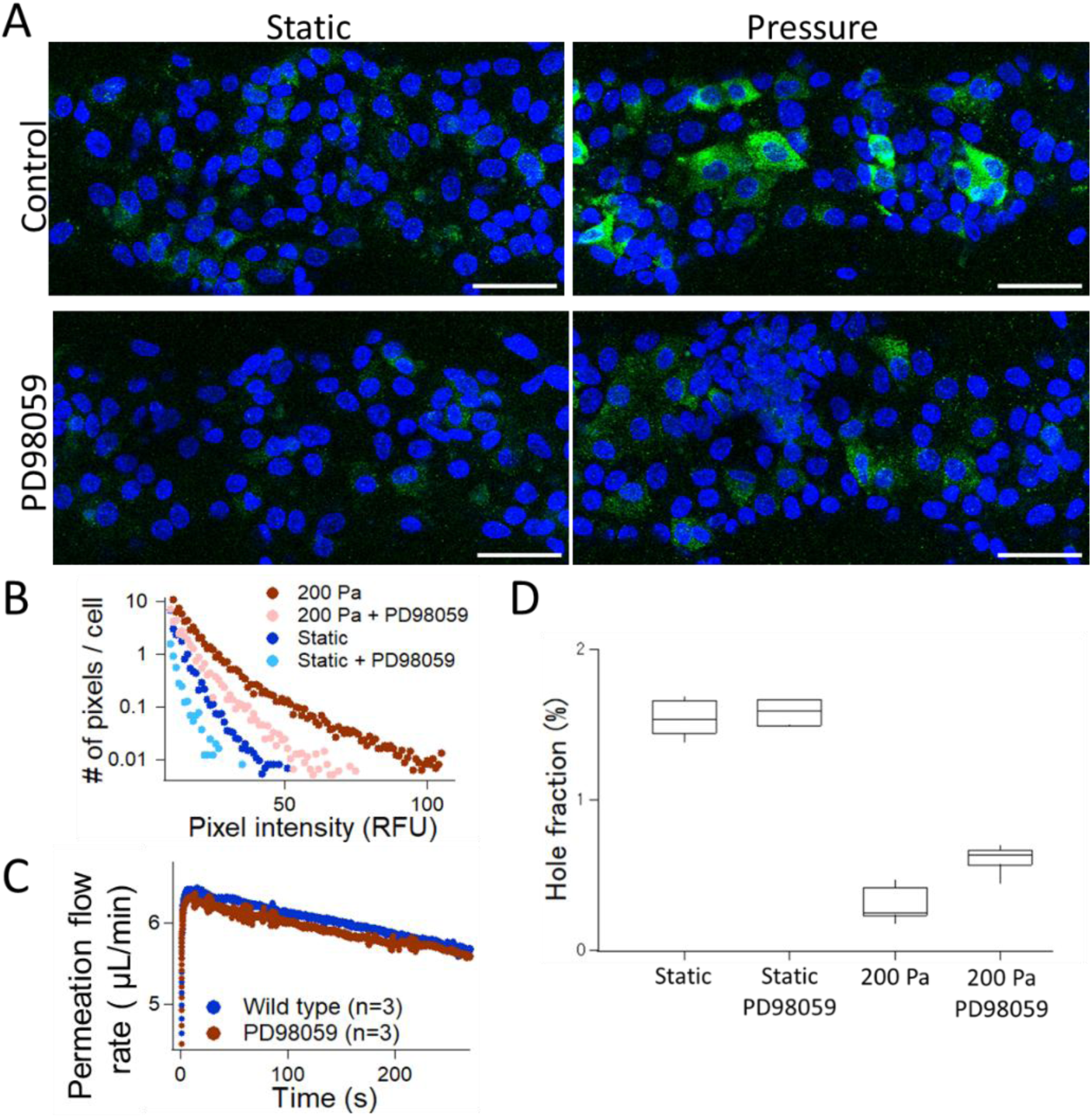
Inhibition of ERK does not interfere with barrier enhancement. **(A)** MIPs of MV in static conditions or treated with PD98059 (upper and lower panels) and placed in static conditions or under pressure; labels: phosphorylated ERK in green and DNA in blue. The scale bars correspond to 100 µm. **(B)** Intensity distribution of the pixels in the MIPs of panel (D) divided by the number of nuclei in the image. **(C)** Averaged permeation flow rate over three MVs in each condition as a function of time for an intraluminal pressure of 200 Pa. **(D)** Hole surface fraction on MV in static vs. pressure conditions using or not the MEK1/2 inhibitor PD98059.

**Supplementary Fig. S6:**
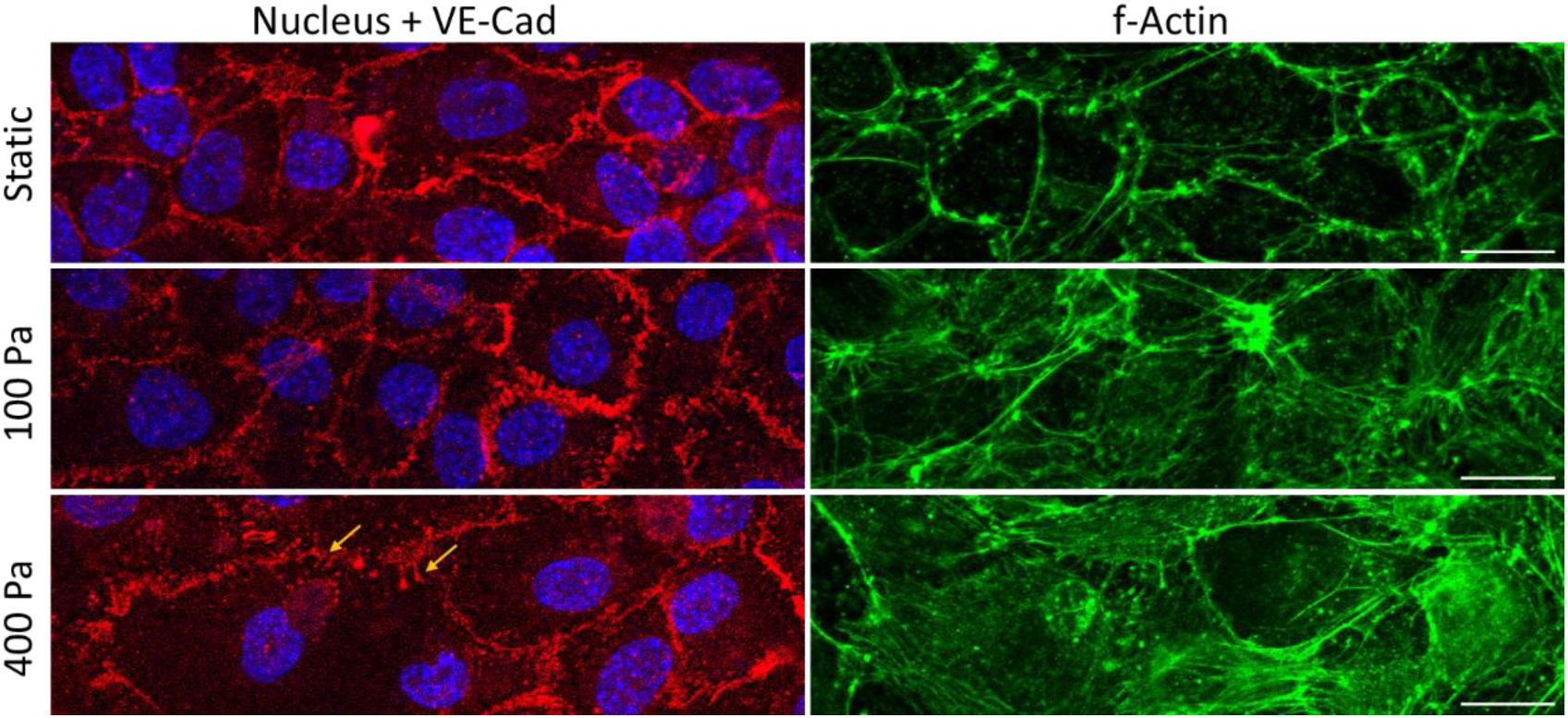
MIP of VE-Cad in static or after pressure stimulation with 1 µM Y-27632. MIPs in static conditions and after the application of 100 or 400 Pa in the same imaging conditions as in main text; labels: f-Actin in green, DNA in blue, and VE-Cad in red. The orange arrows indicate cell protrusions. The scale bars correspond to 20 µm.

## Notes

### Competing Interest Statement

The authors have declared no competing interest.

