## Supplementary Information for "Intraluminal pressure triggers a rapid and persistent reinforcement of endothelial barriers"

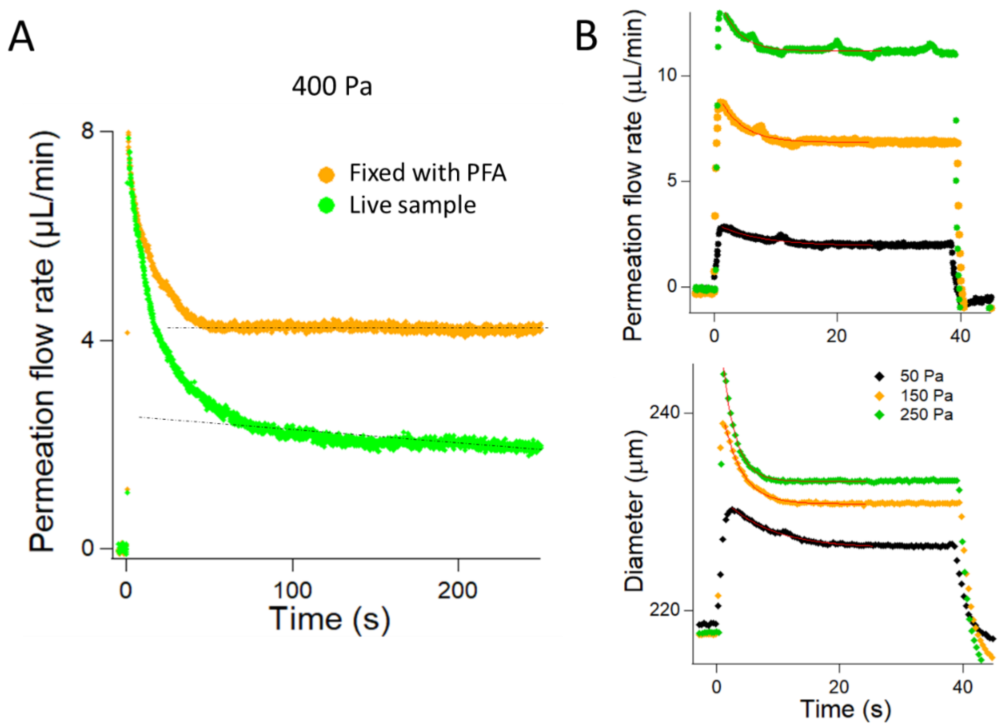


**Supplementary Figure S1:** *Characterization of the permeation flow and MV deformation.* **(A)** Permeation flow rate as a function of time for the same MV live or fixed. The initial value of the permeation is comparable, and the reinforcement of the barrier is lost after fixation. **(B)** The flow rate through the collagen slab is shown in the upper panel for two different intraluminal pressure stimulation of 50 and 250 Pa. The deformation of the tube is represented in the lower panel. The transient regimes of deformation and permeation are closely correlated, as evidenced by the mono-exponential fitting of the deformation relaxation rates, which are 7.7 ± 0.4, 3.03 ± 0.09, and 2.13 ± 0.04 s, compared to the corresponding flow rate relaxation rates of 7.1 ± 0.4, 3.70 ± 0.15, and 2.70 ± 0.08 s for intraluminal pressure stimulations of 50, 150, and 250 Pa, respectively. This correlation arises from the deformation of the tube being linked to a transient regime of additional fluid injection, commonly referred to as the capacitance effect in microfluidics (*50*).

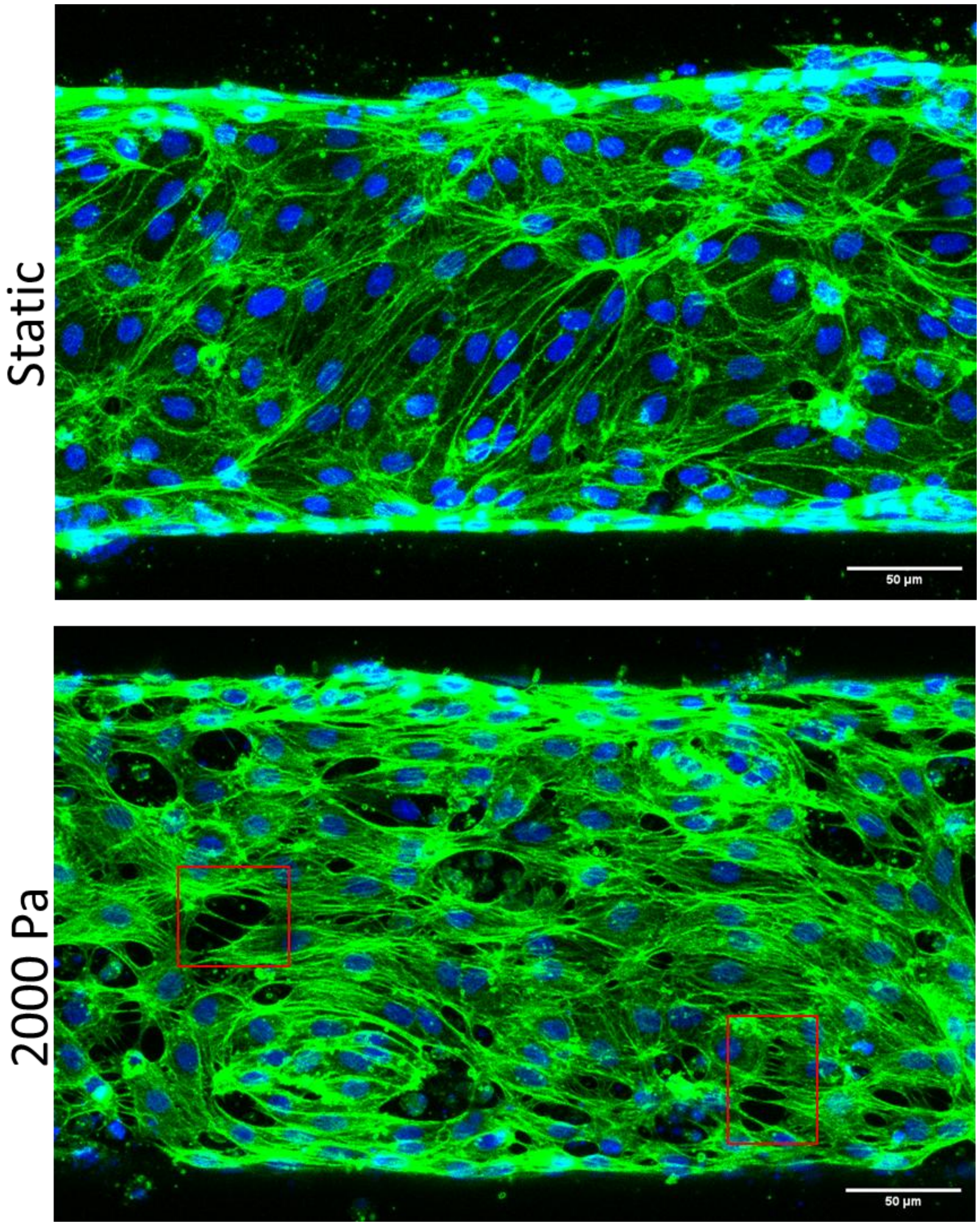


**Supplementary Fig. S2**: *Structural inspection of MVs after stimulation with an intraluminal pressure of 2000 Pa*. The upper panel is the MIP in static conditions and the lower panel presents the stressed MVs. The red rectangles present paracellular holes in the tissue. labels: f-Actin in green, DNA in blue. The scale bars correspond to 50 µm.

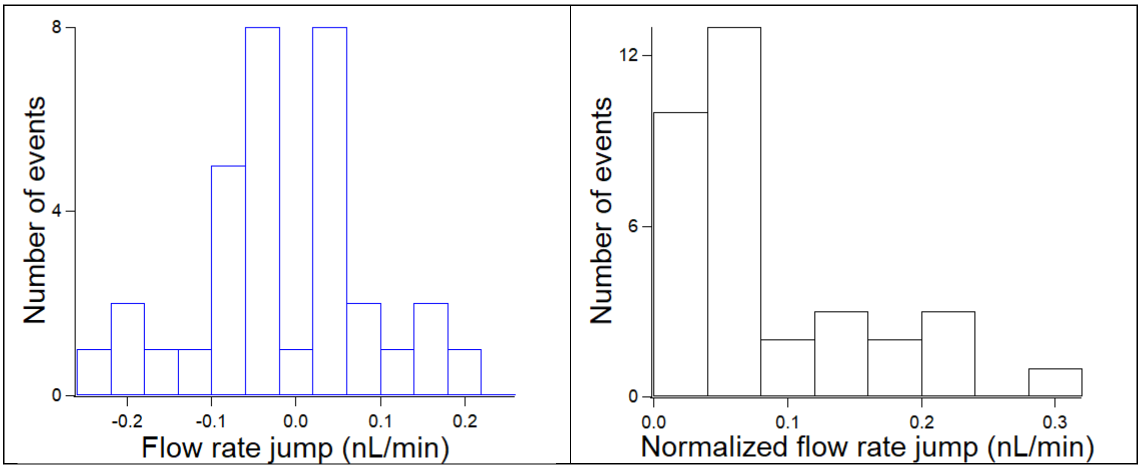


**Supplementary Fig. S3**: *Analysis of the flow rate jumps at 1500 Pa*. The left panel shows the histogram of the flow rate jumps. Note that the probability of positive and negative jumps is roughly equal. The histogram at the right is the same data after normalizing the jumps.

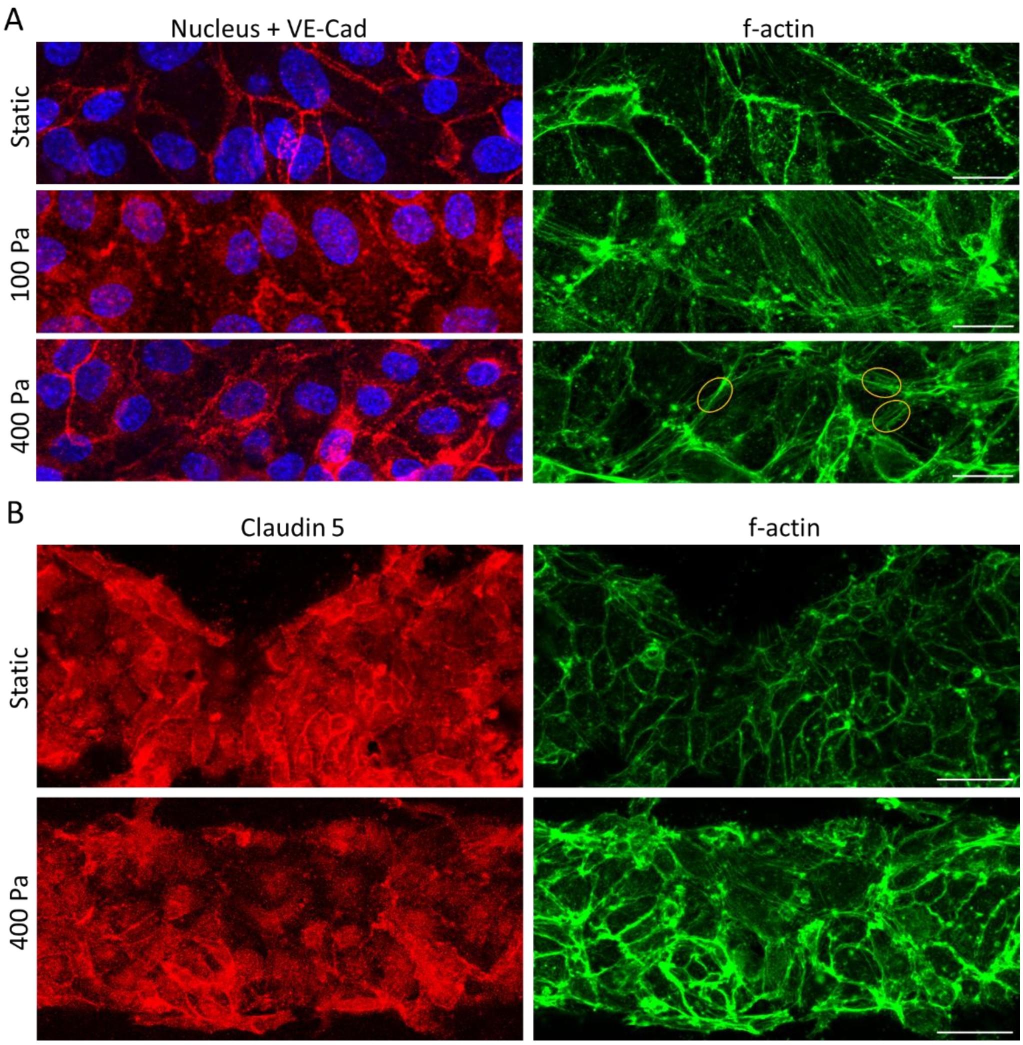


**Supplementary Fig. S4**: *MIP of VE-Cad and Claudin in static or after pressure stimulation.* **(A)** MIPs in static conditions and after the application of 100 or 400 Pa in the same imaging conditions as in main text; labels: f-Actin in green, DNA in blue, and VE-Cad in red. The scale bars correspond to 20 µm. **(B)** MIPs in static conditions and after the application of 400 Pa using a pixel size of 0.313 µm and a vertical inter-frame interval of 2.5 µm; labels: f-Actin in green, DNA in blue, and Claudin in red. The scale bars correspond to 50 µm.

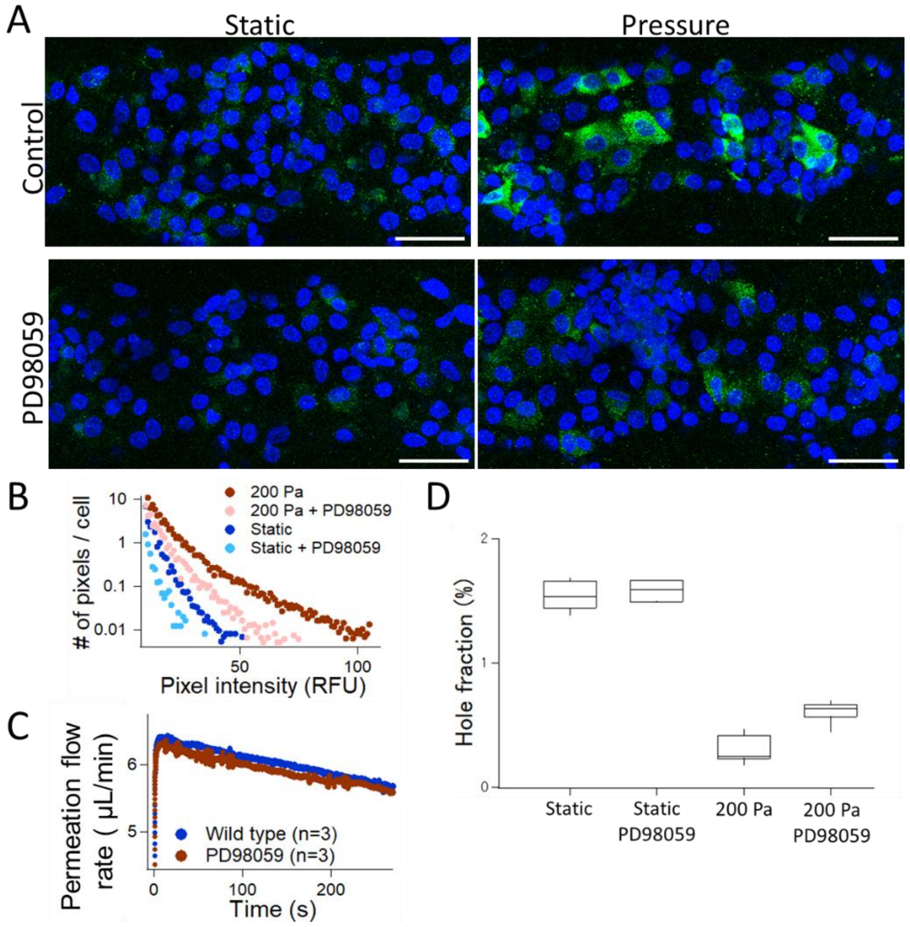


**Supplementary Fig. S5:** *Inhibition of ERK does not interfere with barrier enhancement.* **(A)** MIPs of MV in static conditions or treated with PD98059 (upper and lower panels) and placed in static conditions or under pressure; labels: phosphorylated ERK in green and DNA in blue. The scale bars correspond to 100 µm. **(B)** Intensity distribution of the pixels in the MIPs of panel (D) divided by the number of nuclei in the image. **(C)** Averaged permeation flow rate over three MVs in each condition as a function of time for an intraluminal pressure of 200 Pa. **(D)** Hole surface fraction on MV in static vs. pressure conditions using or not the MEK1/2 inhibitor PD98059.

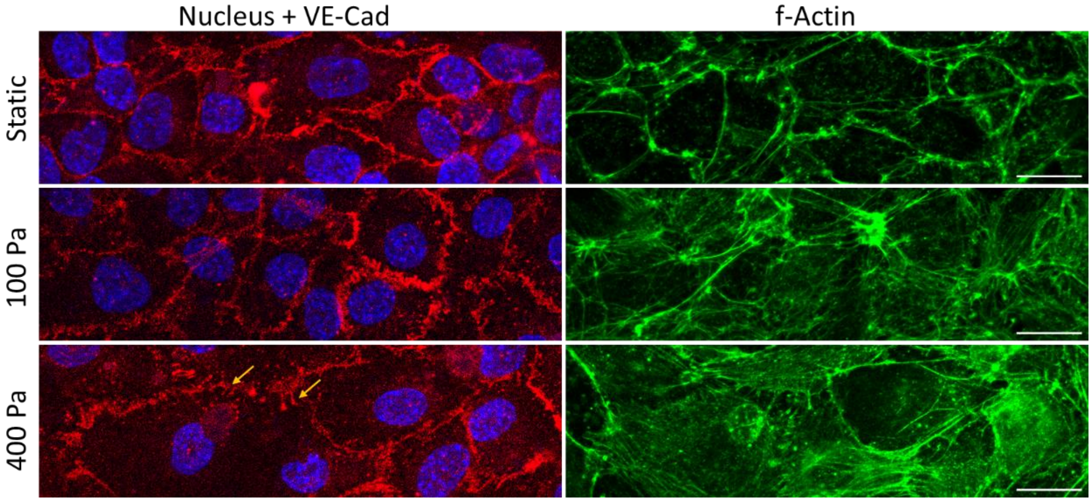


**Supplementary Fig. S6:** *MIP of VE-Cad in static or after pressure stimulation with 1 µM Y-27632.* MIPs in static conditions and after the application of 100 or 400 Pa in the same imaging conditions as in main text; labels: f-Actin in green, DNA in blue, and VE-Cad in red. The orange arrows indicate cell protrusions. The scale bars correspond to 20 µm.
